# Using eDNA to elucidate Silver and Bighead Carp range expansion in two Missouri River tributaries in eastern South Dakota

**DOI:** 10.1101/2023.11.15.567277

**Authors:** Lindsey A. P. LaBrie, Jeff S. Wesner, Hugh B. Britten

## Abstract

**Objective:** This study used environmental DNA (eDNA) sampling methods to determine the possibility of Bighead Carp *Hypophthalmichthys nobilis,* and Silver Carp *H. molitrix*, range expansion above barriers to fish movement in two tributaries of the Missouri River in eastern South Dakota.

**Methods:** We collected water samples above and below two perceived barriers to fish movement: a natural chain of waterfalls in the Big Sioux River and a spillway at the downstream end of a manmade reservoir in the Vermillion River. We used filtration methods to collect invasive carp eDNA from water samples and implemented qPCR techniques to quantify the amount of eDNA in each sample. A total of 213 samples and 38 field blanks were collected over two years (2021/2022).

**Result:** We detected carp eDNA below both barriers in both years, and above the barrier in the Vermillion River in 2022. The probability of detecting eDNA below the barrier in the Big Sioux River was 51% (95% Credible Interval (CrI): 2 to 97%) and 77% (95% CrI: 8 to 99%) in the Vermillion River. The probability of detecting eDNA above both barriers was significantly smaller: 1% (95% CrI: 0.02 to 24%) for both rivers.

**Conclusion:** The detection of positive samples above the spillway barrier in the Vermillion River provides the first evidence that Bighead and Silver Carp may have expanded their range to habitats upstream of their documented range in eastern South Dakota.

**Impact Statement:** This study demonstrates the utility of using eDNA sampling methods to detect Bighead and Silver Carp in areas of both known and unknown invasive carp presence in smaller tributary streams to the Missouri River.

## Introduction

Bighead Carp *Hypophthalmichthys nobilis,* and Silver Carp *H. molitrix*, were introduced into the southern United States in the early 1970s as a means of biocontrol of algae in aquaculture ponds and wastewater treatment facilities (Kolar et al. 2005). Soon after their introduction, both species escaped and began establishing populations throughout the Mississippi River basin and beyond (Nico et al. 2023a, 2023b). In Midwestern rivers, these species have become a threat to many freshwater lotic systems since their arrival in the early to mid 1990s (Kolar et al. 2005) due to their ability to outcompete native planktivorous fishes and alter ecosystems (Freedman et al. 2012; Pendleton et al. 2017; Phelps et al. 2017). In the Missouri River Basin, many barriers limit the upstream movement of both invasive and native fishes. The barriers include dams on the mainstem Missouri and low-head dams and impoundments in the James, Vermillion, and Big Sioux Rivers (Figure 1). Gavin’s Point Dam is a major barrier that prevents upstream movement of these carp into upstream sections of the Missouri River, but they may freely swim between the mainstem Missouri and its tributaries in the stretch of river below Gavin’s Point Dam.

**Figure 1:**
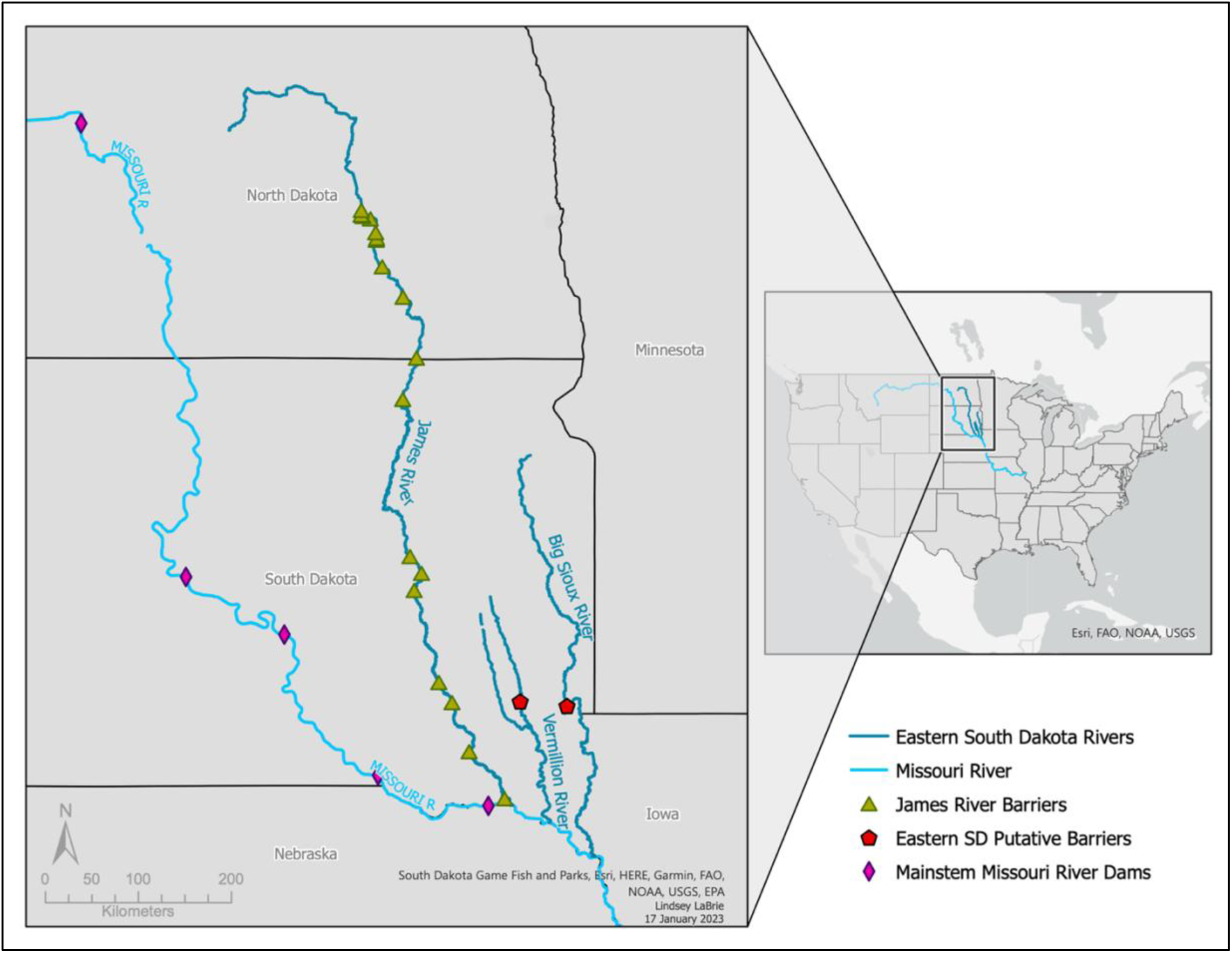
A map of barriers to upstream fish movement in the Missouri, James, Vermillion, and Big Sioux Rivers. Dams on the mainstem Missouri River are shown as purple diamonds, poor barriers to fish movement such as low-head dams and rock impoundments in the James River are shown as green triangles (not all barriers are shown at the current map scale), and the putative barriers in the Big Sioux and Vermillion Rivers are shown as red pentagons.

Streams in eastern South Dakota are on the northwestern-most leading edge of the Bighead and Silver Carp invasion (Hayer et al. 2014). Bighead and Silver Carp first officially appeared in South Dakota in 1998 and in 2003 respectively (Kolar et al. 2005). In the spring of 2019, above-average snowfall followed by rapid melting and early spring rains caused major flooding events throughout rivers in the Midwest (Bagwell and Peters 2019; Flanagan et al. 2020). In eastern South Dakota, this flooding impacted multiple tributaries to the Missouri River (Francis and Melliger 2021), causing them to rise to record-high flood stages. The flooding allowed potential pathways for the spread of invasive carp above two perceived barriers to upstream fish movement: a chain of natural waterfalls in Sioux Falls in the Big Sioux River and a man-made spillway at the downstream end of Lake Vermillion in the Vermillion River. In fact, carp of an undetermined species were visually confirmed swimming over the spillway in the Vermillion River (B. Schall, South Dakota Department of Game, Fish, and Parks, personal communication) immediately after it was compromised during the flooding in 2019. The presence of Bighead and Silver Carp above these barriers in the Big Sioux and Vermillion Rivers was previously determined to be nonexistent, but the flooding created new uncertainty surrounding the invasion front in southeastern South Dakota.

Environmental DNA (eDNA) water sampling has been used to detect Bighead and Silver Carp in other systems and has especially targeted barriers to fish passage into the Great Lakes ecosystems (Erickson et al. 2016; Erickson et al. 2017; Jerde et al. 2013; Song et al. 2017), but there is a lack of its implementation in smaller rivers, especially in the Missouri River Basin. eDNA is a beneficial tool to use because physically sampling an organism is not required, therefore reducing operating costs and minimizing disturbance of the ecosystem, while still providing valuable contemporary evidence for the presence of a species (Thomsen et al. 2012). eDNA taken from water samples allows for the detection of organisms in waterbodies where populations are in low or unknown quantities (Lodge et al. 2012). This is especially important when considering invasive species in novel environments (Erickson et al. 2016). Positive eDNA detections of invasive organisms in waterbodies where their status was previously unknown can then trigger management actions to prevent the unchecked spread of the species.

The purpose of this study was to measure the presence and quantity of invasive carp eDNA below and above two major barriers to fish movement in the Big Sioux and Vermillion Rivers (a chain of natural waterfalls and a man-made spillway, respectively) to determine if recent flooding events allowed for Bighead and Silver Carp movement into habitats above those two barriers. Secondary aims were to evaluate the efficacy of our eDNA sampling regime by 1) determining the number of samples needed to obtain a positive eDNA detections, 2) comparing the effects of filtering method on the amount of eDNA in each sample, 3) comparing the effect of two different extraction protocols on the overall amount of eDNA in each sample, and 4) determining the overall probability of detecting Bighead and Silver Carp eDNA above and below each barrier in both rivers. Data collected from this study is intended to help inform local and basin-wide management efforts to control invasive carp populations in eastern South Dakota and beyond.

## Methods

### Study organisms and sites

The Big Sioux River is a fifth-order stream that begins in Roberts County, South Dakota and flows 674 river kilometers southward through South Dakota to its confluence with the Missouri River on the South Dakota-Iowa border (Omernik 1987). It is the most populated river basin in South Dakota (Dieterman and Berry 1999). There exists a major barrier to upstream fish movement in Sioux Falls—a chain of natural waterfalls called Falls Park. The Vermillion River is a fifth-order stream that flows 242 river kilometers from the confluence of the East and West Fork Vermillion Rivers southward through South Dakota to its confluence with the Missouri River on the South Dakota-Nebraska border (Braaten and Berry Jr 1997). A large spillway (12.5m high) at the downstream end of Lake Vermillion in the East Fork Vermillion River acts as a potential barrier to upstream fish movement (Hayer et al. 2014). Both the Big Sioux River and Vermillion River experience intermittent patterns of flooding and drought, and flows are highly dependent on the amount of precipitation, including snowmelt, and agricultural runoff (Underhill, 1959).

### Sampling methods

Water sampling took place over two field sampling seasons. In the first field season, at each sample location, two water samples were collected using two, 2-Liter high-density polyethylene (HDPE) bottles. In the second field season, one, 2L sample was taken at each sample location. Prior to sampling, all bottles were sterilized with 20% bleach solution for ≥ 10 seconds and rinsed with distilled water (Coulter et al. 2019). Disposable nitrile gloves were worn during sample collection and were replaced after each sampling location. The most downstream location in each river was sampled first, and each successive sample was collected upstream of the previous sample to prevent eDNA cross-contamination. All samples were collected while standing on the shoreline to reduce the risk of boots or waders contaminating the water with invasive carp eDNA. One negative control sample (one, 2-L bottle of distilled water) was collected above and below each barrier. After sampling, all bottles were placed in coolers on ice until filtration was carried out no more than 8 hours after sample collection.

The water samples were filtered through 1.5 μm glass microfiber filters (Cytiva Whatman Binder-Free Glass Microfiber Filters; grade 934-AH circles; 47 mm outside diameter; borosilicate glass) (Eichmiller et al. 2014) using a magnetic polyphenylsulfone filter funnel (Eichmiller et al. 2014; Nukazawa et al. 2018; Coulter et al. 2019). All supplies were sterilized with 20% bleach for ≥ 10 seconds, rinsed with distilled water, and dried with a new paper towel between samples. Water samples were drawn through the filter until the pores were clogged and water could no longer pass through the filter, and the volume of water that passed through was recorded. Each filter was stored in a separate 95% ethanol in a 15 mL polypropylene Falcon tube at -20°C until eDNA extraction could occur.

### eDNA extraction

Prior to extraction, filters were cut in half, with one half returned to the 15 mL Falcon tube in the freezer and the other half cut into three smaller pieces. The pieces were placed in 1.5 mL microcentrifuge tubes and allowed to air dry in a fume hood over a period of ≥ 4 hours until all ethanol had evaporated. Tools were sterilized with 20% bleach for ≥ 10 seconds, rinsed with distilled water, and then rinsed with nanopure water to prevent contamination between filters. eDNA extraction was performed in a designated sequencing room at the University of South Dakota and according to the Qiagen DNeasy Blood and Tissue Extraction protocol (Qiagen 2023) with one modification at the end step to centrifuge 100 μL of elution buffer through the spin columns twice instead of 200 μL once (i.e., “Kristie” protocol (Schmidt et al. 2021)). Extracted samples were stored in 1.5 mL or 2 mL microcentrifuge tubes at -20 °C.

### Quantitative Polymerase Chain Reaction (qPCR) analysis

The real-time quantitative polymerase chain reaction (qPCR) analysis was conducted using an Applied Biosystems™ QuantStudio™ 3 Real-Time PCR System thermocycler (Thermo Fisher). The forward primer, reverse primer, and gBlock used in this study were developed by Erickson et al. (2017) and targeted the mitochondrial gene *nd5*, where the primer sequence binds to an area of the mitochondrial genome that is conserved between both Bighead and Silver Carp, and the probe sequence binds to an area of the mitochondrial genome that is specific to either species (Erickson et al. 2017). Each well had a volume of 20 μL, containing 7 μL molecular grade water, 10 μL TaqMan™ Fast Advanced Master Mix (Thermo Fisher), 1 μL of 10 nanomole forward and reverse primer mix, 1 μL of 5 nanomole probe mix, and 1 μL of extracted eDNA sample. A 313-base pair Megamer ssGene Fragment, or gBlock, containing DNA sequences conserved between both Bighead and Silver Carp was used to create a standard dilution series which served as the eDNA calibration curve for each qPCR run (Erickson et al. 2017). The 96-well plates contained four replicates of each of the following: eight field samples, one field blank, one extraction blank and one positive control. Positive controls were extracted with the DNeasy Qiagen Blood and Tissue Kit from Bighead and Silver Carp tissues. Each plate also contained three spiked replicates of each sample (extracted sample plus 0.1 μL of the 10∧3 dilution per well). Ten, no-template control wells (1 μL nanopure water) and two replicates of a 10∧6 standard dilution series derived from the G-block megamer were included on each plate (Table 1). qPCR cycling conditions followed the protocol in Erickson et al. (2017). We considered each sample to have been positive if at least one sample replicate amplified during qPCR, and negative if none of the replicates amplified.

**Table 1:** A general 96-well plate set up for detecting Bighead and Silver Carp eDNA. “NTC” stands for “no template control.”

|  | 1 | 2 | 3 | 4 | 5 | 6 | 7 | 8 | 9 | 10 | 11 | 12 |
| --- | --- | --- | --- | --- | --- | --- | --- | --- | --- | --- | --- | --- |
| A | Sample 1 | Sample 2 | Sample 3 | Sample 4 | Sample 5 | Sample 6 | Sample 7 | Sample 8 | Field blank | Extraction blank | 10<sup>6</sup> | 10<sup>6</sup> |
| B | Sample 1 | Sample 2 | Sample 3 | Sample 4 | Sample 5 | Sample 6 | Sample 7 | Sample 8 | Field blank | Extraction blank | 10<sup>5</sup> | 10<sup>5</sup> |
| C | Sample 1 | Sample 2 | Sample 3 | Sample 4 | Sample 5 | Sample 6 | Sample 7 | Sample 8 | Field blank | Extraction blank | 10<sup>4</sup> | 10<sup>4</sup> |
| D | Sample 1 | Sample 2 | Sample 3 | Sample 4 | Sample 5 | Sample 6 | Sample 7 | Sample 8 | Field blank | Extraction blank | 10<sup>3</sup> | 10<sup>3</sup> |
| E | NTC | NTC | NTC | NTC | NTC | NTC | NTC | NTC | NTC | NTC | 10<sup>2</sup> | 10<sup>2</sup> |
| F | Spiked sample 1 | Spiked sample 2 | Spiked sample 3 | Spiked sample 4 | Spiked sample 5 | Spiked sample 6 | Spiked sample 7 | Spiked sample 8 | Spiked field blank | Spiked extraction blank | 10<sup>1</sup> | 10<sup>1</sup> |
| G | Spiked sample 1 | Spiked sample 2 | Spiked sample 3 | Spiked sample 4 | Spiked sample 5 | Spiked sample 6 | Spiked sample 7 | Spiked sample 8 | Spiked field blank | Spiked extraction blank | Positive control | Positive control |
| H | Spiked sample 1 | Spiked sample 2 | Spiked sample 3 | Spiked sample 4 | Spiked sample 5 | Spiked sample 6 | Spiked sample 7 | Spiked sample 8 | Spiked field blank | Spiked extraction blank | Positive control | Positive control |

**Table 2:** Average eDNA quantification (copies/μL) for each river, above and below barriers, for each species in both years.

| **2021** |  |  |  |
| --- | --- | --- | --- |
|  |  | Bighead Carp | Silver Carp |
| Vermillion River | Above | 0 | 0 |
|  | Below | 555 | 0.009 |
| Big Sioux River | Above | 0 | 0 |
|  | Below | 18.4 | 1.54 |

| **2022** |  |  |  |
| --- | --- | --- | --- |
|  |  | Bighead Carp | Silver Carp |
| Vermillion River | Above | **0.302** | **0.00960** |
|  | Below | 24.1 | 1.53 |
| Big Sioux River | Above | 0 | 0 |
|  | Below | 2.74 | 0.391 |

### Pilot study

First, to evaluate the number of water sample replicates needed to obtain a positive detection in areas of known Bighead and Silver Carp presence, we took five, 2-L samples in September 2022 at one location in the Big Sioux River (42.495167, -96.477236) and one location in the Vermillion River (42.772914, -96.930374), where the USFWS was contemporaneously performing their eDNA sampling and Dozer trawling for invasive carp. Second, to determine if eDNA quantities varied when following the Qiagen extraction protocol versus the modified extraction protocol (i.e., “Kristie” protocol (Schmidt et al. 2021)), we extracted one half of each filter using the Qiagen protocol and the other half using the “Kristie” protocol (Schmidt et al. 2021) and then analyzed the samples.

### Statistical analysis

We evaluated whether filtering method (i.e., field vs lab) caused a difference in the probability of eDNA detection. We used Program R (version 4.1.1; R Core Team 2022) and the *brms* package (Bürkner 2017) to fit a Bayesian generalized linear mixed model with a Bernoulli likelihood with a logit link with the following model and default priors:

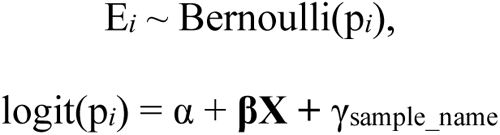

Where E*_i_* is the presence or absence of eDNA in sample *i*, p*_i_* is the probability of detection, α is the intercept, **βX** is a vector of 3 parameters representing two predictors (**X** = filter method, species) and their interaction and γ_sample_name_ is the random intercept for sample name.

To evaluate the probability of detection as a function of species, river, and location in respect to the barriers in each river, we again used a Bernoulli likelihood with a logit link using the following equation:

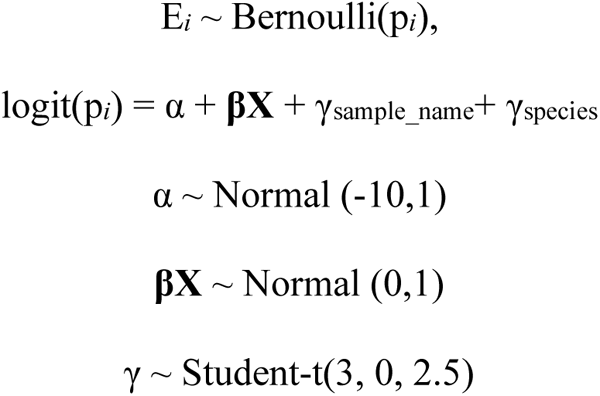

Where E*_i_*, p*_i_*, and α are as defined above, **βX** is a vector of 7 parameters representing three predictors (quantity, location, river) and their interactions, and γ_sample_name_ and γ_species_ are the random intercepts for sample name and species, respectively.

Both models were run using a Hamiltonian Monte Carlo No-U-Turn sampler with 4 Markov chains, 2,000 iterations per chain with a warm-up of 1,000 iterations. We evaluated the fit of the model by running posterior predictive checks and determined it properly converged if Gelman-Rubin statistics were less than 1.1 (Brooks and Gelman 1998). We sampled from the posterior and estimated the relationships for the model and predicted the mean with 95% credible intervals. We ran a sensitivity analysis to assess the impact of the priors on our presence/absence model by doubling the standard deviation values to Normal (-10,2) for the intercept, Student’s t (3, 0, 5) for the random intercepts, and Normal (0,2) for the beta values.

We also visualized how other factors may have affected the quantity of eDNA in our samples using *ggplot2* (Wickham et al. 2016), such as the length of time between sample collection and filtering, sample month, the volume of water filtered, the number of days between filtering and extraction, and the number of days between extraction and qPCR.

## Results

### Water sampling

A total of 213 water samples and 38 field blanks were collected after two years of field sampling. (98 water samples and 20 field blanks in 2021, 115 water samples and 18 field blanks in 2022). In 2021, one-half of the water samples were filtered in the field using a 7.2-liter, manual pump MityVac MIT7400 fluid evacuator (Figure S-1a). The other half of the samples were transported back to the lab and filtered using a 1000 mL Büchner flask (Figure S-1b). In 2022, samples were exclusively filtered using the field filtering method. The average filtered volume was 382 mL (range 100 to 1650 mL). Each year, samples were collected from sampling sites in both rivers three separate times. In 2021, Big Sioux River samples were collected from three below-barrier sites and six above-barrier sites in three sampling sessions. This was expanded to five below-barrier sites and eleven above-barrier sites in 2022 (Figure S2). In 2021, Vermillion River samples were taken at two below-barrier sites and seven above-barrier sites. This was expanded to four below-barrier sites and fourteen above-barrier sites in 2022 (Figure S3).

Invasive carp were observed swimming below the barriers at almost all sampling sessions, except for the below-barrier Vermillion River samples in October and September 2021, where only fish carcasses of various species, including Bighead and Silver Carp, were observed in the water below the spillway. In 2022, we visually observed carp at all water sampling sessions below both barriers.

### qPCR results

Bighead and Silver Carp eDNA was detected below both barriers in 2021 and 2022 in varying quantities (Table 3). In July 2022, Bighead and Silver Carp eDNA was detected in two and three samples respectively above the Lake Vermillion spillway (Figure S4), but not in previous or successive samples. The probability of detecting eDNA below the barrier in the Big Sioux River was 51% (95% Credible Interval (CrI): 2 to 97%) and in the Vermillion River was 77% (95% CrI: 8 to 99%). The probability of detecting eDNA above the barriers was significantly smaller: 1% (95% CrI: 0.02 to 22%) in the Big Sioux River and 1% (95% CrI: 0.02-24%) in the Vermillion River. The probability that detection was higher below the barriers than above the barriers was greater than 99% (Figure 2).

**Figure 2:**
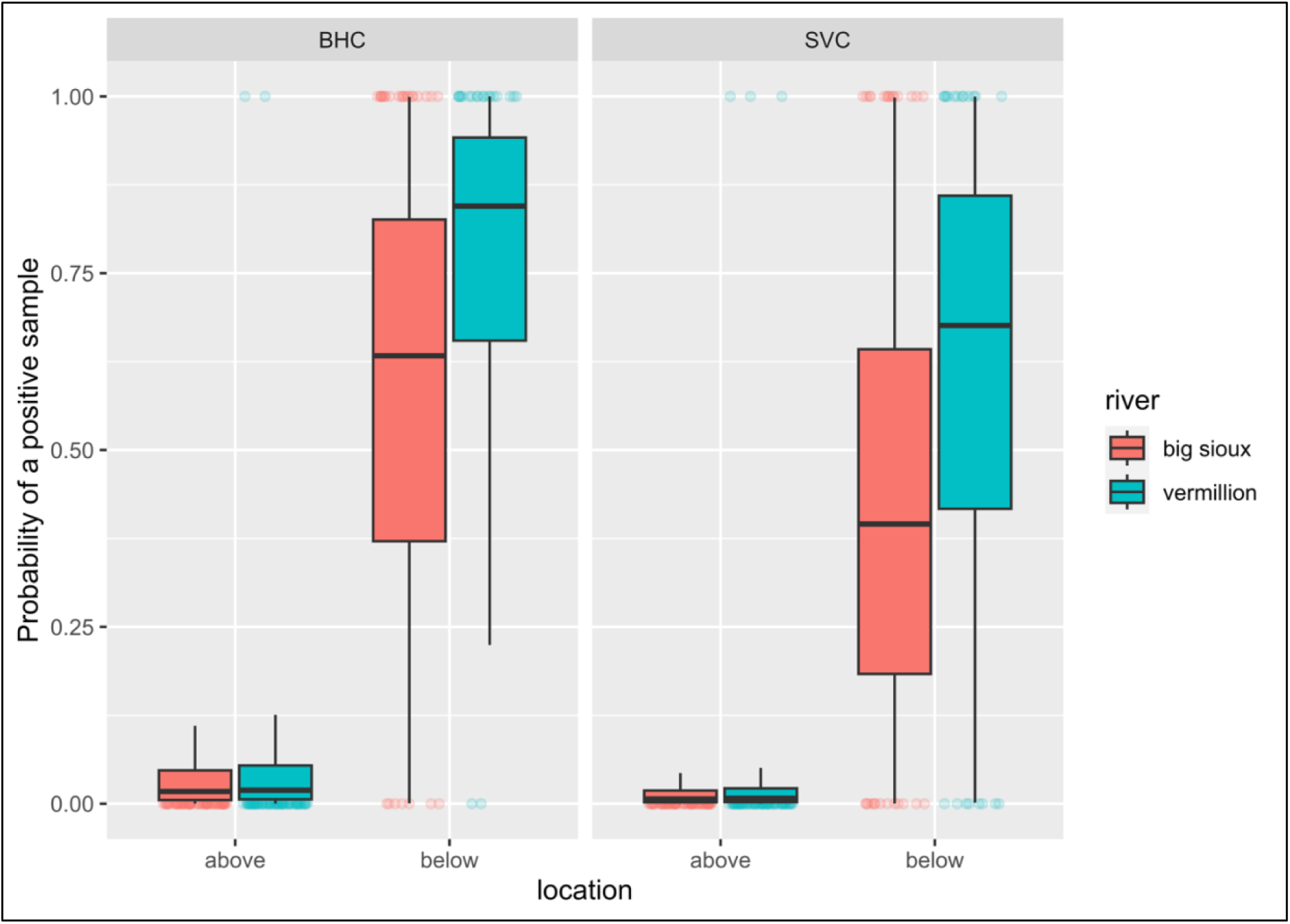
Posterior predictions for the probability of a positive sample for Bighead Carp (BHC) and Silver Carp (SVC) in the Big Sioux and Vermillion Rivers above and below the major barriers to upstream fish movement. The hollow, colored circles represent data points, with zero (0) on the Y-axis indicating no eDNA detection and one (1) on the Y-axis indicating a positive eDNA detection. The probability of a difference between above and below barrier samples was greater than 99%.

There was little difference between the mean quantity of eDNA obtained from field and lab filtering methods, with less than one-hundredth of one percent difference between field and lab filtered samples (0.003%, 95% CrI: 0.001 to 0.6%) (Figure 3). This was the basis for our choice to only perform field filtration in the second sampling season. Through pilot study sampling, we determined that one, 2L water sample at each sampling location was sufficient to obtain a positive detection for both Bighead and Silver Carp in both rivers where eDNA was present, with a 10% probability of false negative detection. The probability of false negatives halved when more samples were taken at each location (i.e., 5% for two samples per location, 2.5% for three samples per location, etc.). The extraction protocol type (i.e.,“Qiagen” versus “Kristie”) appeared to have an impact on mean eDNA quantities, which reinforced our decision to follow the modified extraction protocol described in the methods section above. Other factors such as the length of time between sample collection and filtering, sample month, the volume of water filtered, the number of days between filtering and extraction, and the number of days between extraction and qPCR did not appear to have any strong effect on the amount of eDNA detected with qPCR (Supplementary Figures 5-9).

**Figure 3:**
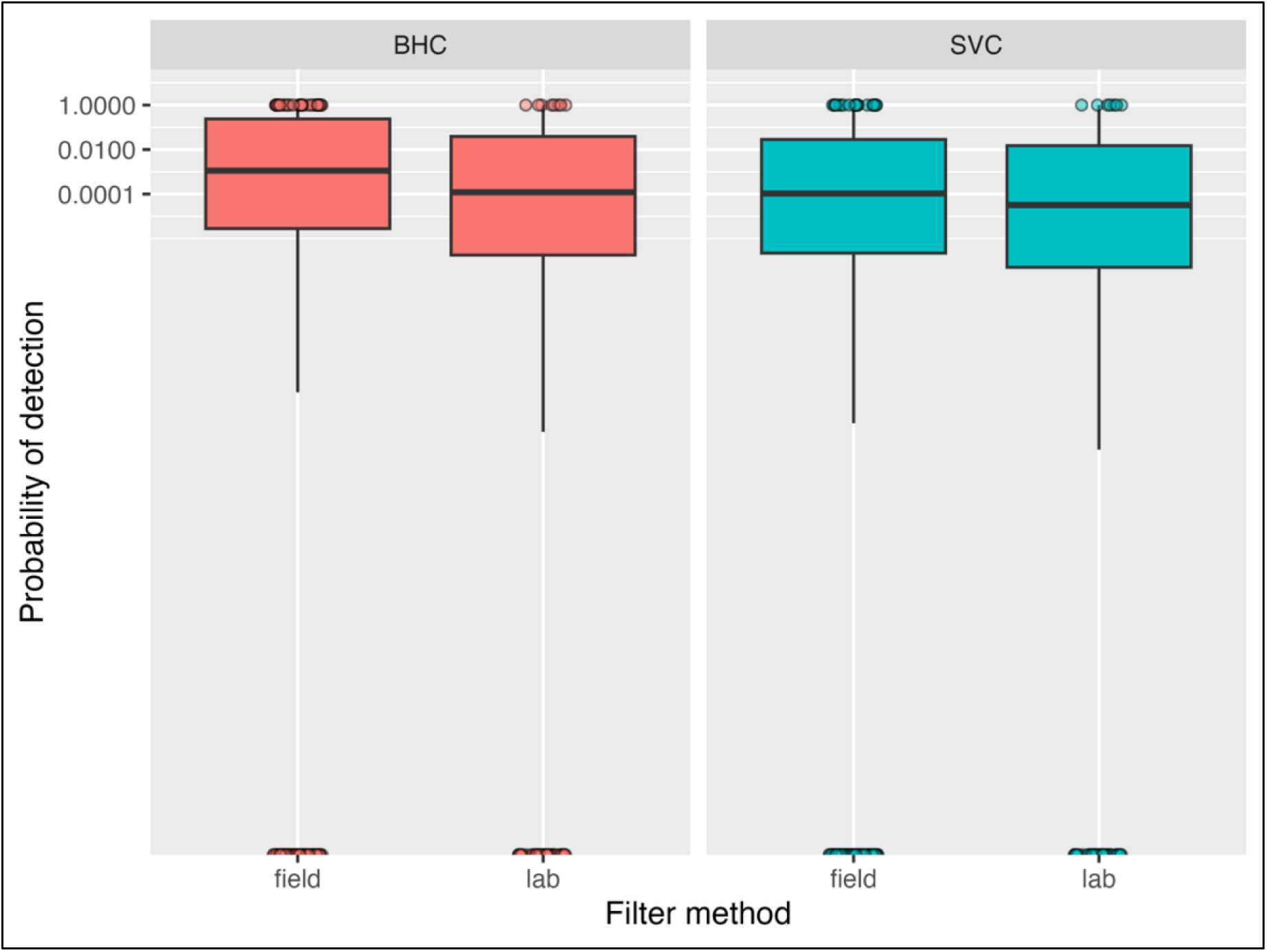
Estimated posterior probability of detecting eDNA from Bighead Carp (BHC) and Silver Carp (SVC) in both rivers (combined) for field and lab filtered samples. The colored circles represent actual data points, with zero (0) on the y-axis indicating no eDNA detection and one (1) on the y-axis indicating a positive eDNA detection. Note that the y-axis is on a log(10) scale.

### Model performance

Both models performed well, and observed data was within the predicted range of the posterior estimates. Trace plots indicated proper mixing of the Markov chains, all R-hat values were < 1.001, and all parameter estimates had large effective sample sizes. The Bayesian R^2^ value for the filter method model was 0.82 (95% credible interval (CrI): 0.73 to 0.89) and for the presence/absence model was 0.76 (95% CrI: 0.67 to 0.84). Posterior predictive checks (Berkhof et al. 2000) generally replicated the shape of the data for both models. The sensitivity analysis showed model parameter estimates were not sensitive to the priors (Figure 4).

**Figure 4:**
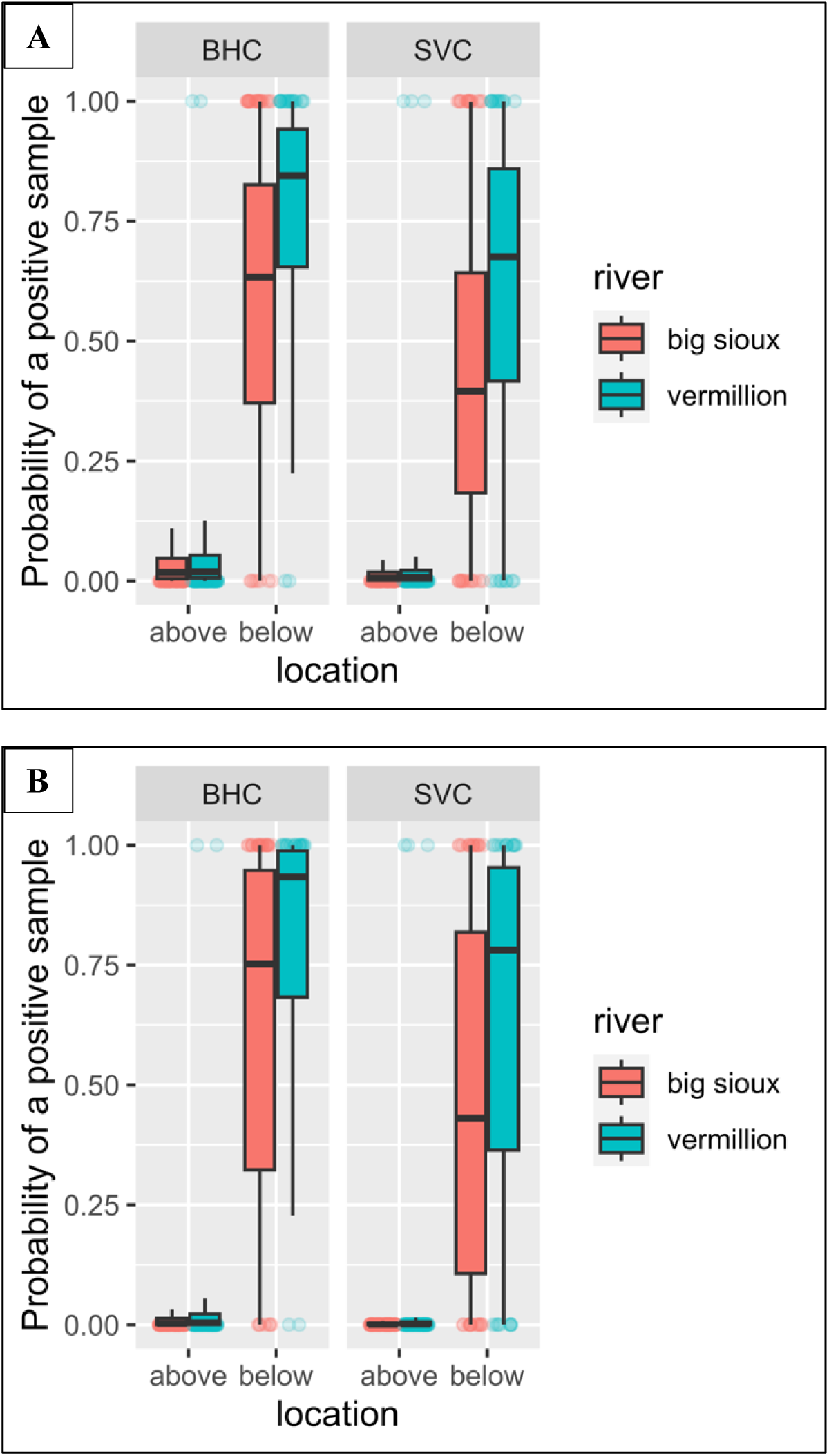
A comparison of the original model estimates (A) and the sensitivity analysis (B) for which the standard deviation of each prior was doubled. The colored circles represent actual data points, with zero (0) on the y-axis indicating no eDNA detection and one (1) on the y-axis indicating a positive eDNA detection.

## Discussion

This study was the first to use eDNA to detect Bighead and Silver Carp above barriers to upstream movement in Missouri River tributaries in eastern South Dakota. We were able to demonstrate the utility of the filtration method to quantify eDNA from water samples within these two rivers. The filtration method used in this study has the potential to be more accessible than other eDNA sampling methods. The manual pneumatic pump and magnetic filter funnel are relatively inexpensive, easily portable, and can be deployed on-site and allow for immediate filtration upon water sample collection. We determined that filtering water from one 2-L sample was sufficient to detect eDNA at least 90% of the time when both species of carp are present. The number of days between filtration and extraction, or the number of days between extraction and qPCR analysis did not appear to affect the quantity of eDNA detected in each positive sample.

The probability of detecting eDNA in our samples below the barriers was high (>54%) and the probability above the barriers was low but greater than zero (<1%) in both rivers. We obtained three positive eDNA detections above the spillway in the Vermillion River, within Lake Vermillion in June 2022, but not in previous or successive samples. We are confident that positive samples above the spillway were true positives because there was no evidence of eDNA contamination in the field blanks or qPCR no-template controls. There have been no official reports of invasive carp above the spillway, however it is possible that invasive carp have been persisting in Lake Vermillion in relatively low numbers and that we simply did not sample in areas where they were present during the other rounds of sampling. The presence of boats on Lake Vermillion during our July 2022 sampling session could explain the positive detections above the spillway barrier. Invasive carp DNA could have sloughed off the hull of a boat that had previously been in carp-positive waters, resulting in positive detections. Another explanation is that invasive carp initiate spawning movements in early summer when the hydrograph begins to rise (Erickson et al. 2016). Because eDNA has short persistence times in freshwater ecosystems (Eichmiller et al. 2014), it is possible that the invasive carp made an upstream spawning run out of the lake and into the Vermillion River immediately prior to sampling. Both could offer potential explanations for the positive samples above the barrier within the lake.

South Dakota is predicted to experience an increase in climate extremes, including persistent severe droughts, at least a 15% increase in annual precipitation, and an increase in the intensity and interval of severe flooding (Frankson et al. 2022). The effects of flooding have been evident in the recent past during the flood of 2019, when Falls Park became inundated with water and the spillway at Lake Vermillion became compromised, thus providing potential pathways for invasive carp to pass barriers to fish movement. Future eDNA water sampling or physical sampling methods (i.e., capture of live individuals via electrofishing) will be required to confirm the presence of live invasive carp above the barriers in both rivers. Water sampling could be expanded to other small tributaries to assess the presence of these fish in finer detail throughout eastern South Dakota. It may also be necessary to eventually reconsider the effectiveness of these barriers during flooding events.

## Conclusion

This study provides evidence that eDNA is a useful technology to implement in smaller tributary systems on the leading edge of the current invasive carp range. We determined that filtering one, 2-L sample per site provided a true positive eDNA detection at least 90% of the time. We established a baseline for fish presence above barriers in the Big Sioux and Vermillion Rivers, however, we recommend continued monitoring to increase the level of certainty of fish presence above the barriers in both rivers, especially in the Vermillion River. We recommend future eDNA water sampling to further elucidate the status of these species above the barriers, coupled with physical sampling efforts to gauge invasion presence and risk into the future.

## Acknowledgements

We thank T Devine and J Larsen for their assistance with water sampling and processing, and K Schmidt for demonstrating the qPCR protocol. Dr. A Coulter provided comments and revisions to the manuscript. Funding for this study was provided South Dakota Game Fish and Parks (Grant number F-16-R-1) and the United States Fish and Wildlife Service (Grant number F20AP11067-00).

## Data Availability Statement

Data and R code used in this manuscript are available in the GitHub repository: https://github.com/lindseylabrie/carp_eDNA

## Ethics Statement

This work was conducted under an approved protocol from the University of South Dakota’s Institutional Animal Care and Use Committee (protocol # 10-08-20-23D).

