## Supplemental Figures for "Using eDNA to elucidate Silver and Bighead Carp range expansion in two Missouri River tributaries in eastern South Dakota"

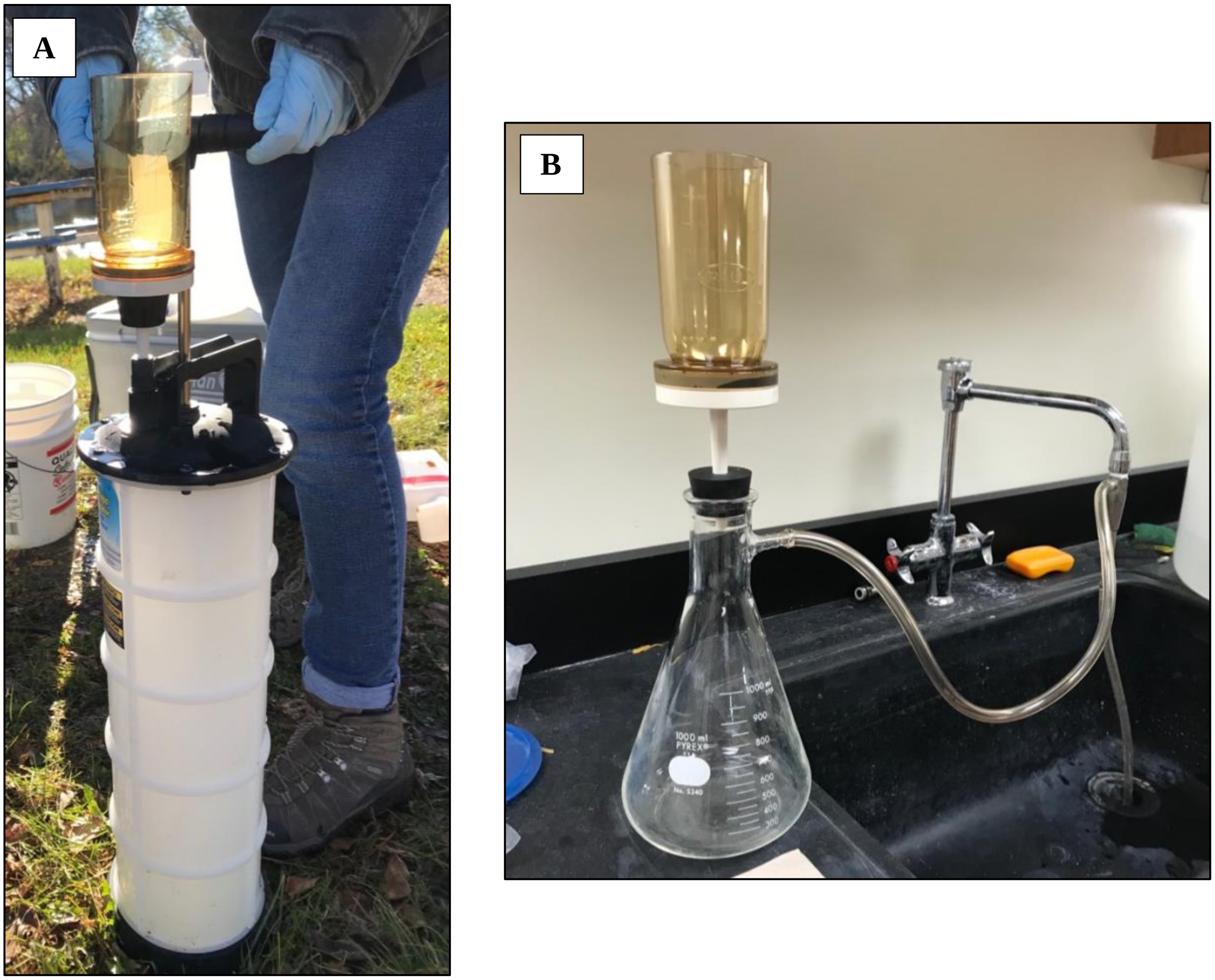


Figure S1a) The MityVac fluid extractor vacuum and magnetic filter funnel setup for eDNA water sample filtering in the field. S1b) The Büchner flask and magnetic filter funnel setup for eDNA water sample filtering in the lab.

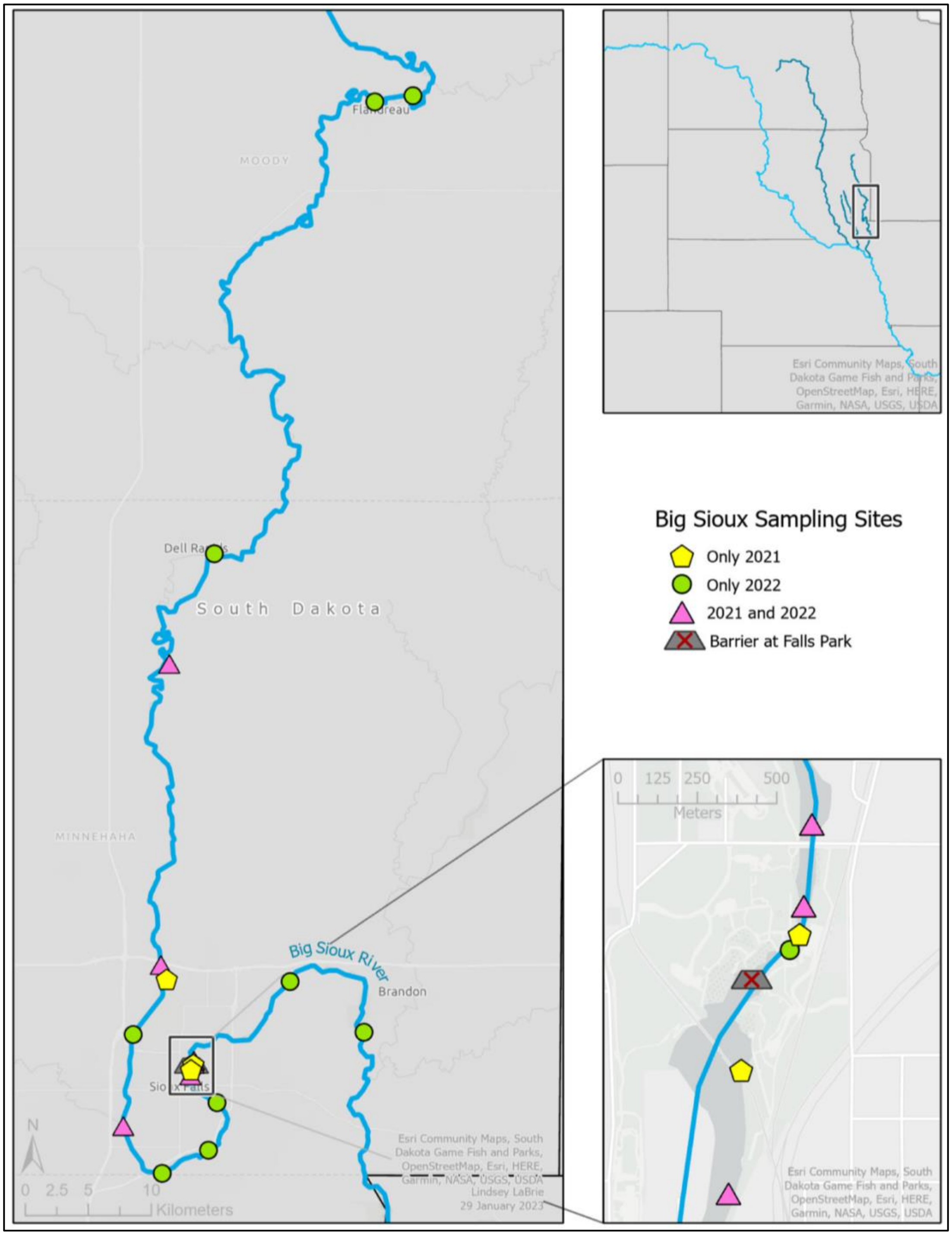


Figure S2) eDNA water sampling locations in the Big Sioux River. The inset map depicts a zoomed in portion of the river below and above Falls Park in Sioux Falls, SD. Yellow pentagons represent areas sampled only in 2021, pink triangles represent areas sampled in both years, green circles represent sampling locations that were added in 2022.

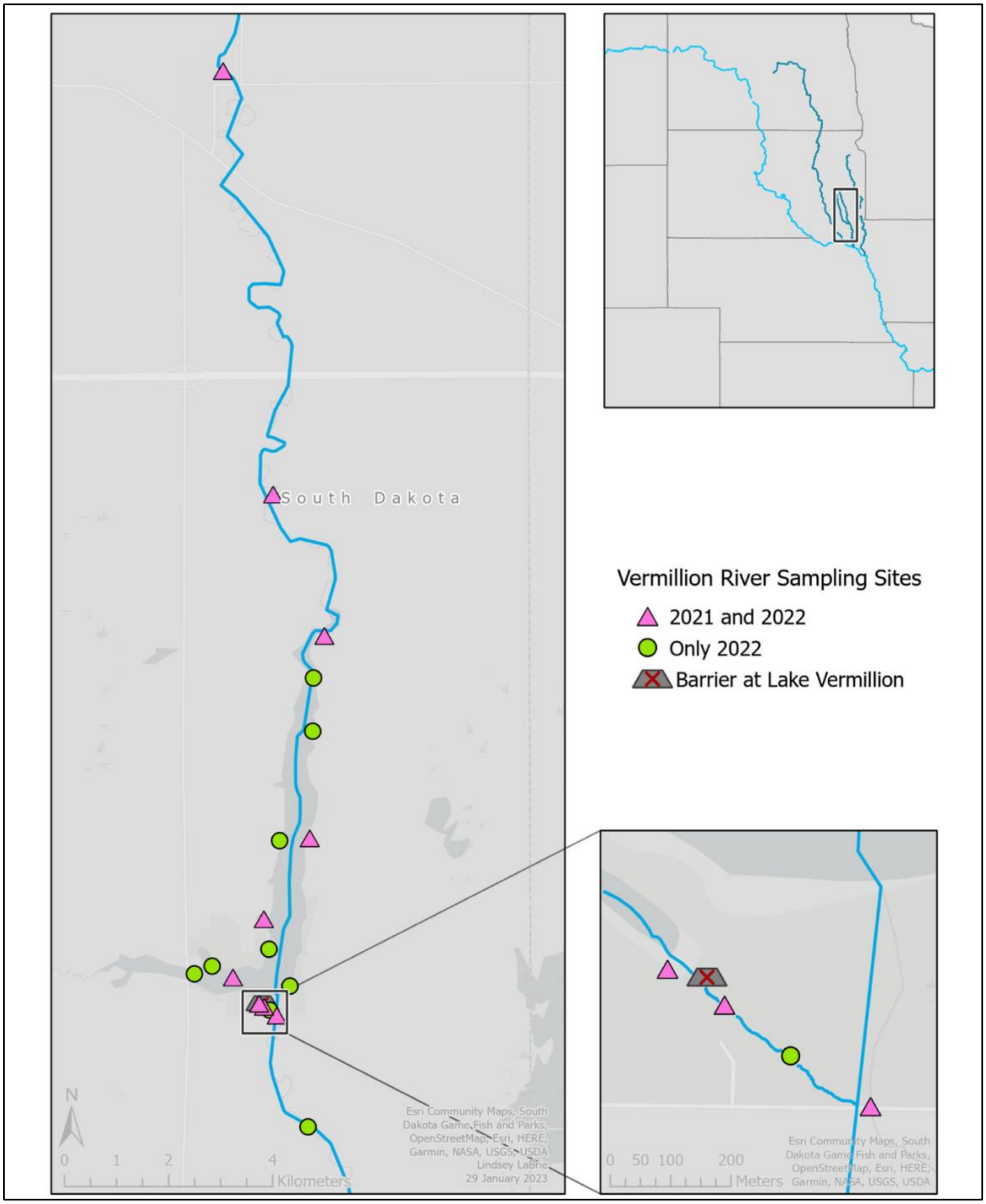


Figure S3) eDNA water sampling locations in the Vermillion River. The inset map on the bottom right depicts a zoomed-in portion of the river below the spillway at Lake Vermillion. Pink triangles represent areas sampled in both years, and green circles represent sample sites that were added in 2022.

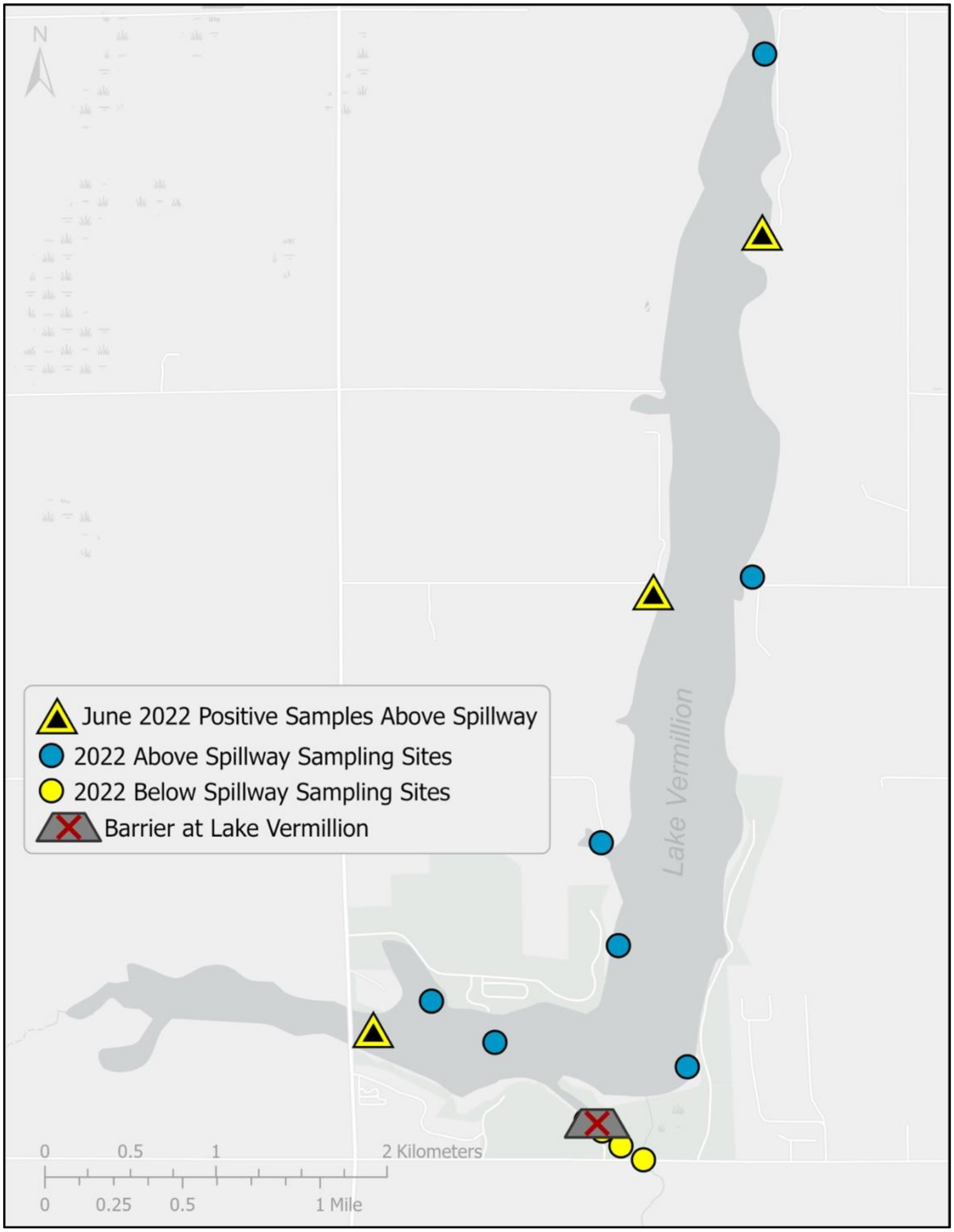


Figure S4) Locations of positive eDNA detections (represented by yellow triangles) in Lake Vermillion. Samples were collected in June 2022.

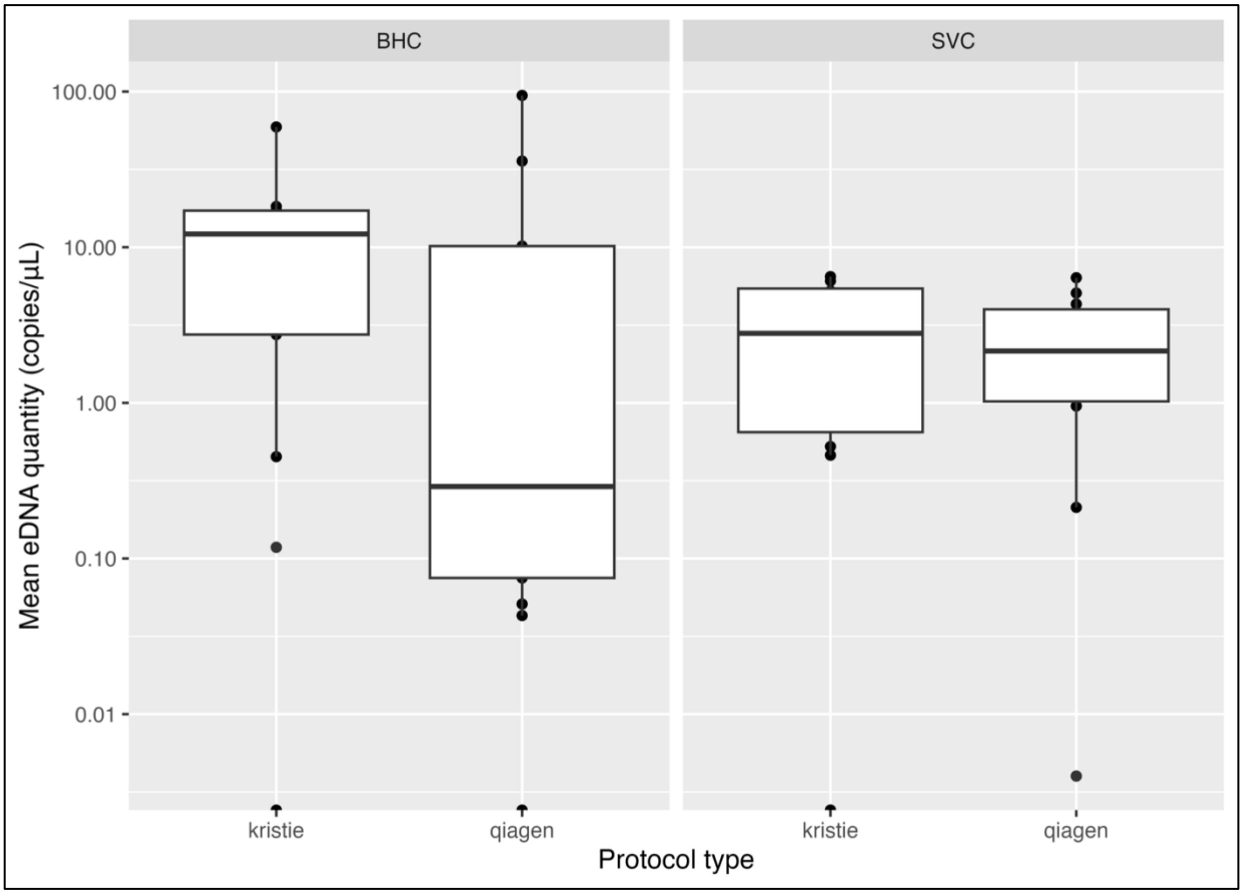


Figure S5) The effect of extraction protocol on mean eDNA quantities (in copies/µL) for detection of both species (Bighead Carp: BHC; Silver Carp: SVC). “Kristie” is the modified protocol with the 2x-100 µL elution buffer step, whereas “Qiagen” is the original 1x-200 µL elution buffer step. Note that the y-axis is on a log(10) scale.

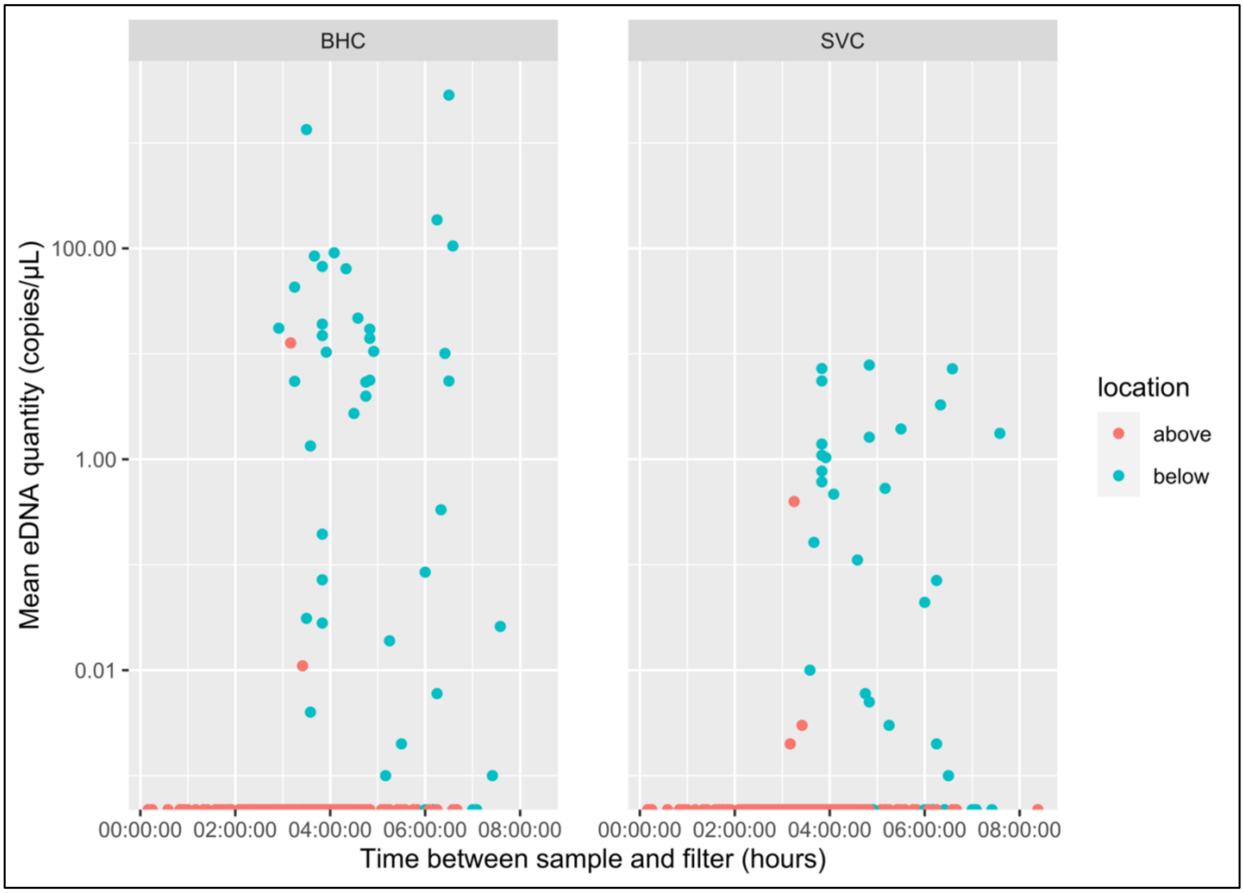


Figure S6) The relationship of the time between sample collection and sample filtration on the quantity of eDNA detected in each sample (in copies/µL), colored by location (above and below the barriers in both rivers). The data is from both field seasons.

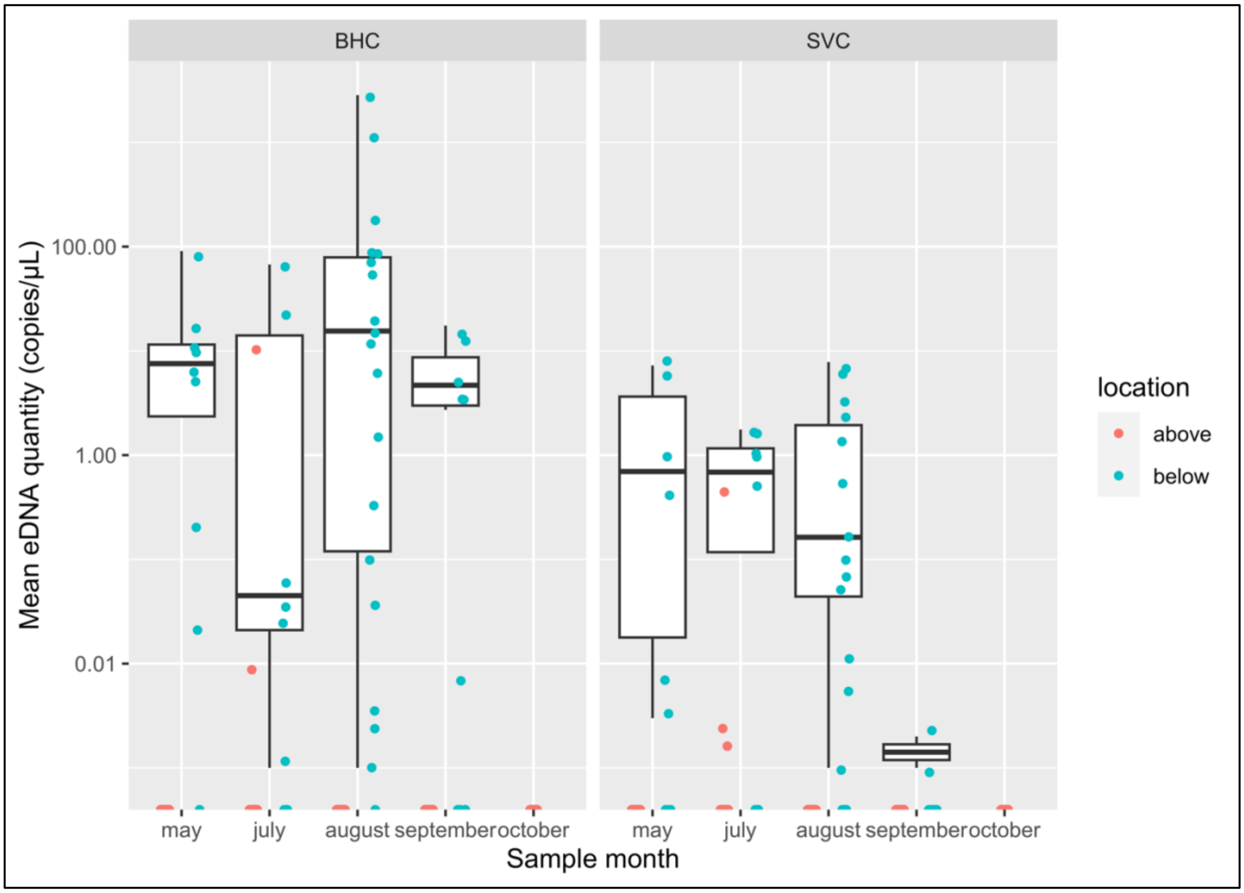


Figure S7) The relationship between mean eDNA quantities (in copies/µL) and sample month. Overall, Bighead Carp (BHC) detections tend to result in higher eDNA quantities than Silver Carp (SVC) detections. The data for samples below the spillway in October are missing, because we were unable to sample below the spillway in October 2021. The data is from both field seasons.

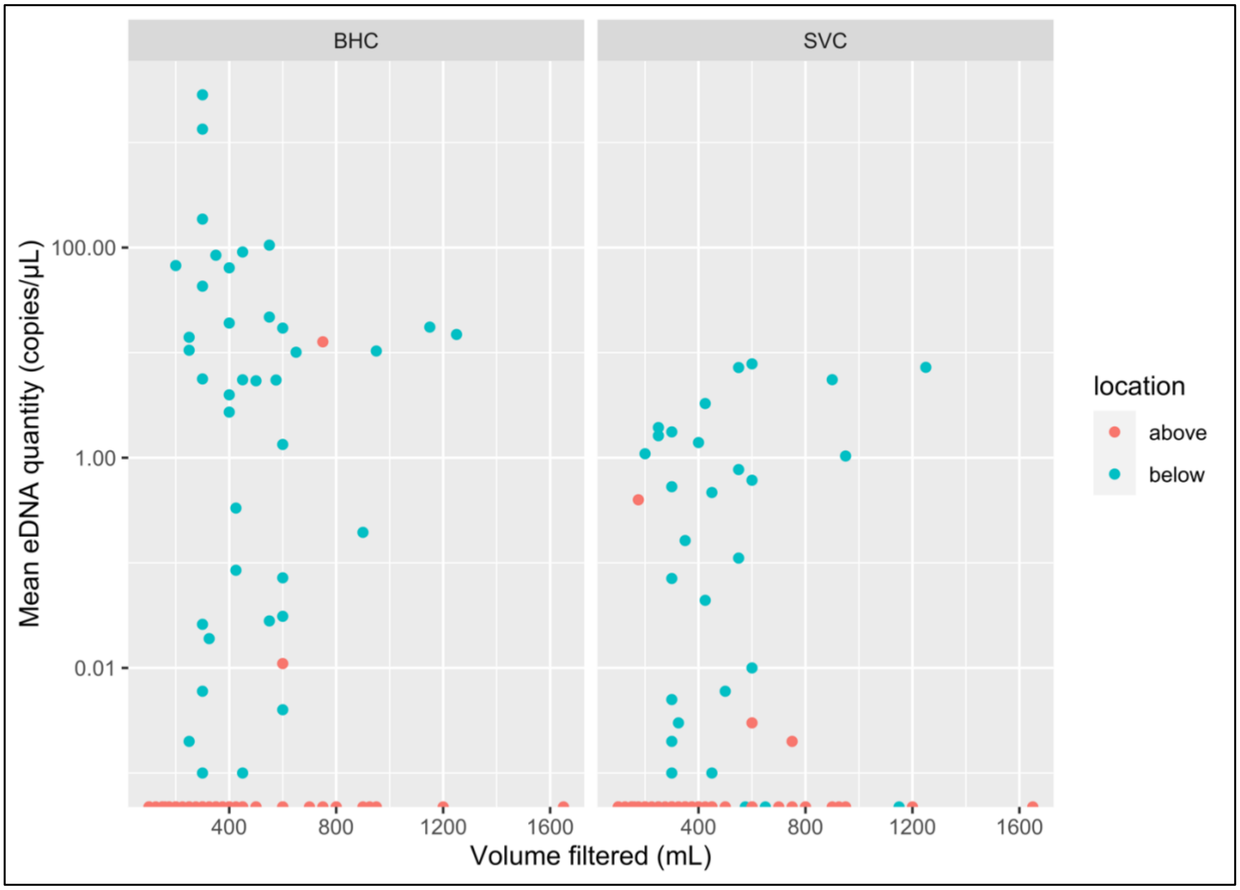


Figure S8) The relationship between volume of water vacuumed through the filters and the average eDNA quantity (in copies/μL) for each species (Bighead Carp: BHC; Silver Carp: SVC). The data is from both field seasons.

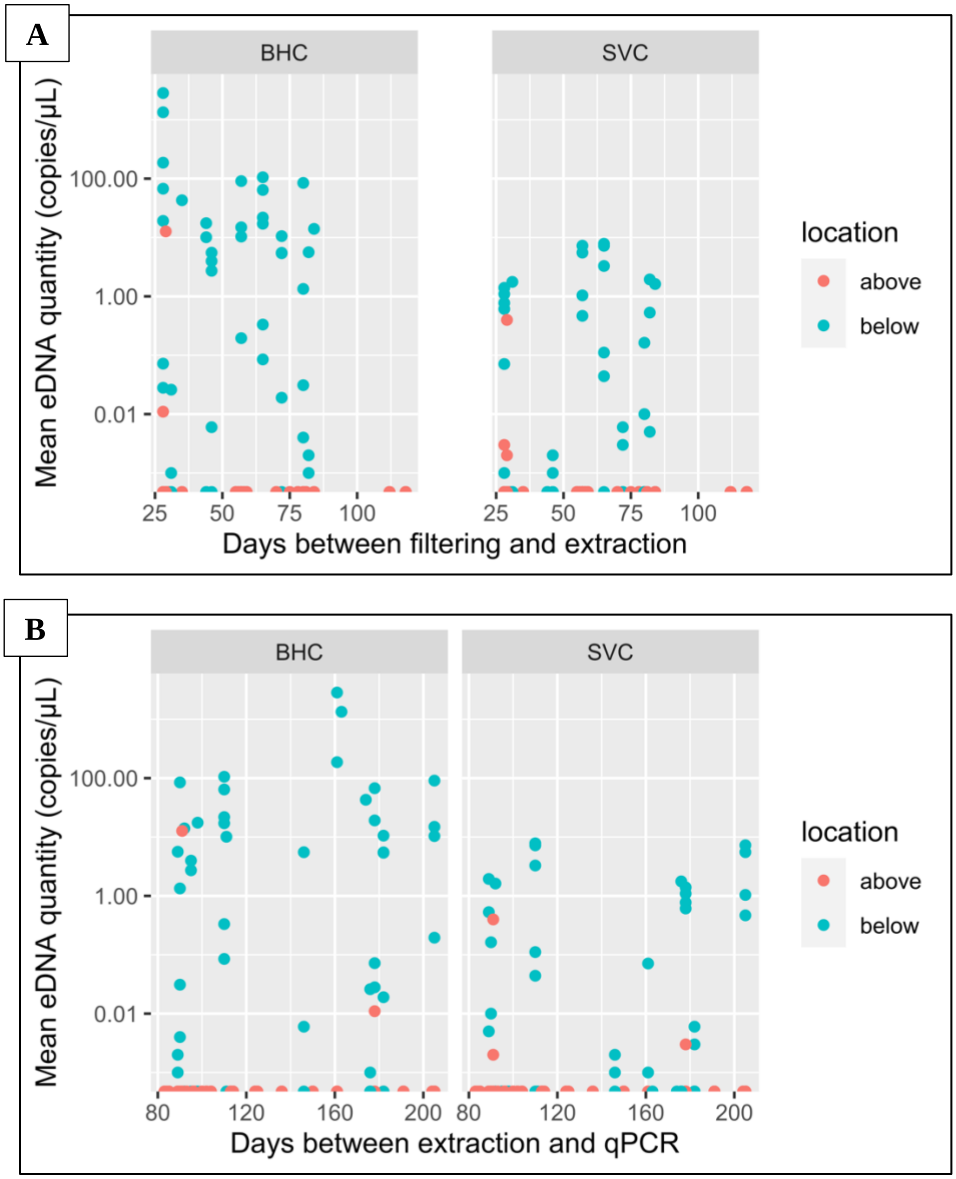


Figure S9a) The relationship between the number of days between filtering and extraction on mean eDNA quantity (in copies/µL) for Bighead Carp (BHC) and Silver Carp (SVC). S9b) The relationship between the number of days between extraction and qPCR on mean eDNA quantity (in copies/µL) for Bighead Carp (BHC) and Silver Carp (SVC).
